# Annotation of structural variants with reported allele frequencies and related metrics from multiple datasets using SVAFotate

**DOI:** 10.1101/2022.06.09.495527

**Authors:** Thomas J. Nicholas, Michael J. Cormier, Aaron R. Quinlan

## Abstract

**Background:** Identification of impactful genetic variants from DNA sequencing data relies on increasingly detailed filtering strategies to isolate the small subset of variants that are more likely to underlie a disease phenotype. Datasets reflecting population allele frequencies of different types of variants have been demonstrated as powerful filtering tools, especially in the context of rare disease analysis. While such population-scale allele frequency datasets now exist for structural variants (SVs), it remains a challenge to match SV calls between multiple datasets and thereby correctly estimate the population allele frequency of a putative SV.

**Results:** We introduce SVAFotate, a software tool for SV matching that enables the annotation of SVs with variant allele frequency and related information. These annotations are derived from known SV datasets which are incorporated by SVAFotate. As a result, VCF files annotated by SVAFotate offer a variety of annotations to aid in the stratification of SVs as common or rare in the broader human population.

**Conclusions:** Here we demonstrate the use of SVAFotate in the classification of SVs with regards to their population frequency and illustrate how annotations provided by SVAFotate can be used to filter and prioritize SVs. Lastly, we detail how best to utilize these SV annotations in the analysis of genetic variation in studies of rare disease.

## Background

Structural variants (SVs) encompass a diverse range of genomic changes that vary considerably in type and size, but are commonly defined as any DNA variant that consists of at least 50 nucleotides (1). SVs can alter DNA copy number or structure, impact gene dosage, and contribute to human phenotypes (2–4). Accurate identification of SVs is challenging as many SVs arise in paralogous and repetitive regions, resulting in inconsistencies between samples not only in terms of the presence or absence of a given SV, but also in the predicted breakpoints of the event. It is estimated that the average human genome harbors at least 8,000 SVs, encompassing millions of bases of DNA sequence, when compared to the human reference genome (5). However, when considering individual Mendelian phenotypes, including human diseases, the vast majority of these SVs are likely inconsequential. Thus, the identification of causal or otherwise impactful SVs relies on increasingly sophisticated SV variant filtering and prioritization pipelines.

Traditional genomic annotation tools, like the Variant Effect Predictor (6) or SnpEff (7), rely on identifying overlaps with known genomic features to provide annotations that aid in identifying SVs that may have phenotypic consequence. However, these methods are generally more applicable to other variant types including single nucleotide variants (SNVs) and insertions-deletions (INDELs). Annotation tools have been created specifically for the complexities of SVs (8–10) and may add useful information to help define SVs of interest. Existing software tools attempt to prioritize SV calls with scores that reflect the potential pathogenicity of a given SV call (11–14). However, many of these tools are trained on or incorporate previously published data and few consider the entirety of available SV-specific datasets. Furthermore they have limited capacity to integrate additional SV callsets as they continue to be generated.

Large-scale reference datasets of human genetic variation have enabled the measurement of accurate allele frequencies (AFs) and provide the opportunity to stratify variants as common or rare in the general population (15,16). Prioritizing variants based on their population frequency provides an effective prioritization strategy for SNVs and INDELs, especially in the context of rare disease. Filtering based on AF substantially reduces the number of putative genetic variants for analysis, especially when combined with additional annotations and expected inheritance patterns (17).

Until recently, population-scale measures of SV allele frequencies have been limited. The Database of Genomic Variants (18) represents the first human SV-specific reference dataset, consisting of SV calls from a variety of studies, methods, and platforms. In recent years, however, collaborations involving the whole-genome sequencing (WGS) of thousands of human samples have generated extensive SV datasets making population-level AFs and related metrics available for SV analysis. These include the CCDG (19), gnomAD (20), and 1000G (21) SV datasets, all of which were generated using different samples, sequencing protocols, and SV calling methods. Unsurprisingly, given the distinct methodologies used to create each dataset and human population growth (22), the vast majority of all identified SVs are rare (average of 85% with AF < 0.01) and many SVs are limited to only a few samples, or are “singletons” (i.e., found in one individual, **Supplementary Figure 1**).

By focusing on overlapping genomic coordinates to identify SVs observed in multiple datasets, we find that the fraction of SVs of the same variant type (SVTYPE) shared by multiple datasets decreases as the degree of overlap required among them increases (**Figure 1**).

**Figure 1.**
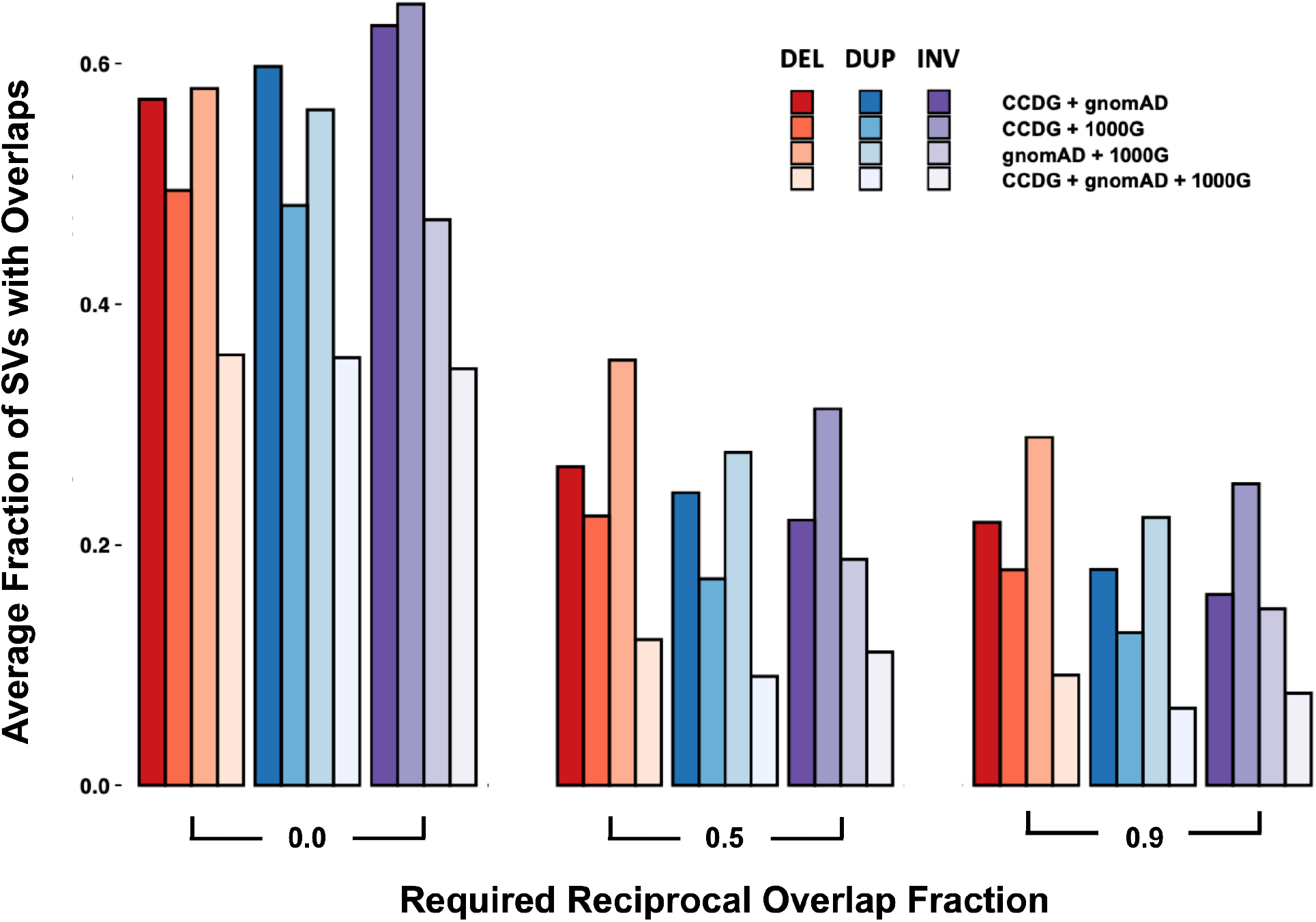
Matching SVs from different datasets based on shared SVTYPE and genomic overlaps. The average fraction of overlaps between deletions (DEL, in red), duplications (DUP, in blue), and inversions (INV, in purple) from CCDG, gnomAD, and 1000G are identified using varying amounts of required reciprocal overlap. Higher required reciprocal overlap fractions correspond to more exact genomic coordinate matches. Each dataset is compared to one another (CCDG + gnomAD, CCDG + 1000G, and gnomAD + 1000G) and overlaps with a different required reciprocal fraction are calculated. The fraction of total SVs found to have overlaps given the required reciprocal overlap fraction is found for each respective dataset and the average of these fractions is plotted. Finally, the average fraction of SVs found to have overlaps in all datasets (CCDG + gnomAD + 1000G) is found for each SVTYPE and at each required reciprocal overlap fraction.

This observation highlights two important aspects when comparing SV calls across datasets. First, since the vast majority of genetic variants in the human population are rare, most SV calls found in these datasets are unique to those collections, and therefore, no single dataset sufficiently represents the full spectrum of potential SVs in the general population. Second, as opposed to SNV and INDEL variant calls, matching SV calls across datasets varies depending on both the presence or absence of the variant as well as the required amount of overlap between potential matches. This additional “spatial uncertainty” illustrates the possible variability in genomic coordinates between calls, but also the difficulty in determining whether SV calls with overlapping coordinates represent the same or different variants (23). These observations are not surprising given the differences in samples, sequencing, and variant calling between datasets, but differences in read length, read depth, and insert size will also result in differing coordinates for the same SV. Thus, trying to match observed SVs to SVs in these population datasets is a challenge that requires special consideration, especially in the context of rare disease.

Recognizing this obstacle, we have created SVAFotate as a tool that provides the means to aggregate SV calls from multiple SV population datasets and create summaries of AF-relevant data into simple annotations that are added to SV calls based on default or user-determined SV matching criteria. Primarily, this enables the classification of SVs within a VCF (24) file as being either common, rare, or unique to an individual dataset with respect to the thousands of samples in published datasets. Being able to differentiate SVs based on their population frequency enables powerful filtering strategies in the context of SVs for rare disease analysis. Here, we describe the functionality of SVAFotate, demonstrate the effectiveness of its annotations for filtering SV calls, and describe recommendations for using SVAFotate in rare disease analyses.

## Implementation

SVAFotate is a Python-based command line tool that annotates an input VCF file with allele frequencies and related information from SVs reported in population-scale datasets. Two distinct file types are required as input for SVAFotate: an input SV VCF file and an input BED (25) file containing known or reported SV calls with accompanying AF information.

### Input SV VCF File

SVAFotate has been tested on VCFs created from various SV callers and is compatible with any VCF that includes SVTYPE (preferably END and SVLEN also) in the INFO field. All SV calls in the VCF are internally converted into a BED format for the purposes of identifying overlapping genomic coordinates with the SVs provided by the input BED file, and the output from SVAFotate is returned as an annotated VCF file.

### Input BED file

The motivation behind SVAFotate was to enable the comparison of unannotated SV calls and known SVs with computed AFs from multiple population datasets, resulting in SV calls with AF-related annotations. As a result, variants can be analyzed and prioritized based on these population frequency annotations. With that in mind, SVAFotate requires that known population SV calls with pertinent AF data be provided as an input file in the BED format. Provisional BED files corresponding to the GRCh37 and GRCh38 genome reference builds have been created by parsing and compiling SVs and their associated AF data from the CCDG, gnomAD, and 1000G datasets and can be found here: https://github.com/fakedrtom/SVAFotate/tree/master/supporting_data/. The CCDG and 1000G published datasets were generated using the GRCh38 reference and thus for the provided GRCh37 BED file, coordinates pertaining to these datasets were converted to GRCh37 coordinates using the UCSC liftover executable (https://genome.ucsc.edu/cgi-bin/hgLiftOver). Similarly, the gnomAD dataset, which was generated using the GRCh37 reference, was converted to GRCh38 for inclusion in the provided GRCh38 BED file. These BED files define genomic breakpoints of reported SVs, and include additional columns detailing the allele frequency information for each listed SV. The origin of each SV in these files, with respect to the dataset that includes it, is listed and labeled as the SV’s source. Given that each published dataset provides different scores, measurements, and data, in the event that certain data is unavailable from a given dataset, those columns are still included in the BED file with a “NA”. Researchers can also provide custom BED files from their own cohorts, as long as the structure of the file matches SVAFotate’s built-in BED file structure (**Supplementary Table 1**).

### Identifying SV matches

SVAFotate attempts to identify matches for each SV from the input VCF with SVs in the BED file based on congruent SVTYPEs and overlapping genomic coordinates (**Figure 2a**). By default, SVAFotate considers any amount of overlap as sufficient for matching purposes, but allows for a recommended, optional parameter which requires overlapping SVs with the same SVTYPE to share a minimum reciprocal overlap for consideration as a match (see SVAFotate Best Practices). In this manner, SVAFotate allows users to control the specificity with which matching SVs are defined where higher reciprocal overlaps reflect more exact coordinate agreement between potential matches. For each matching SV, corresponding AF metrics from the input BED are saved with the maximum values across all matches being returned, creating annotations that reflect the observed maximum AF (`Max_AF`) and other complementary annotations (**Figure 2b, Supplementary Table 2**). Any individual SV from the input VCF that does not have any matches with SVs from the input BED file, and is therefore unique with respect to those provided SVs, is still annotated by SVAFotate, but as appropriate these annotations will have a value of `0`.

**Figure 2.**
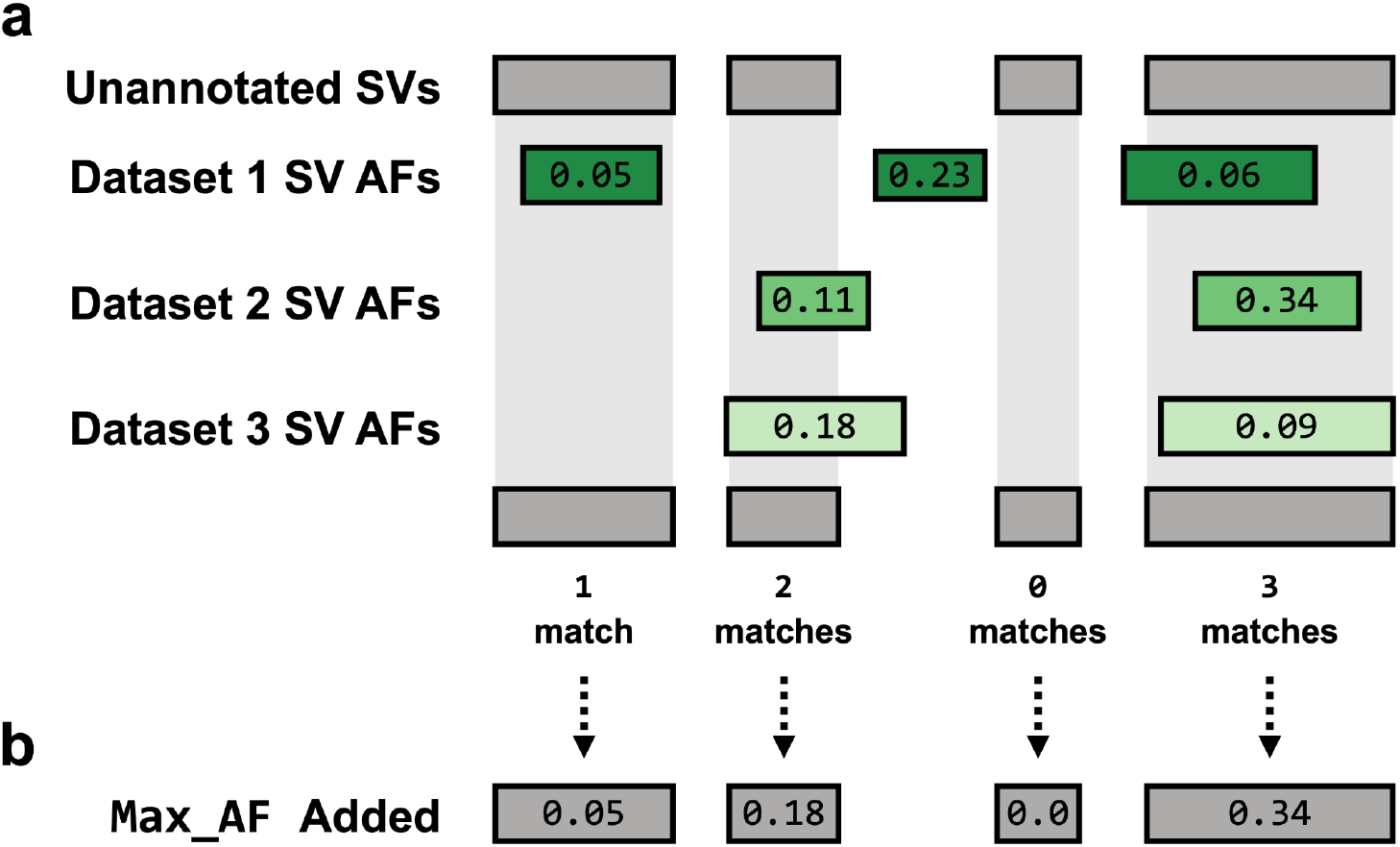
Matching SVs for Annotation Creation. **a**. SVAFotate expects two distinct input files: an unannotated SV VCF file and a BED file which may contain SV calls from multiple population datasets and their accompanying AF metrics. To represent the SV calls in these files, unannotated SVs are illustrated as gray rectangles while SVs from three different datasets, such as CCDG, gnomAD, and 1000G, are represented by green rectangles with their reported population AF included as labels. For this example we will assume that all rectangles represent SVs of the same SVTYPE. SVAFotate attempts to identify matches between unannotated SVs and the SVs present in the BED file by identifying genomic coordinate overlaps that meet user-defined criteria between SVs of the same SVTYPE. Multiple matches are possible, and all AF related data is saved for each match. **b**. SVAFotate is capable of generating multiple annotations that are added to the original VCF file and are each derived using information saved from matching the SVs. The types and variety of annotations added to the VCF are determined by input parameters provided at the command line, but here the example annotation added is the `Max_AF` (default) annotation.

Identifying matching SVs as described is more straightforward for many SVTYPEs, such as, deletions (DELs), duplications (DUPs), and inversion (INVs), but can be more complicated for other SVTYPEs. Insertions (INSs), for example, are often reported as a single base pair genomic coordinate with an accompanying SVLEN that reflects the size of the insertion. SVAFotate still matches INSs based on overlapping coordinates and shared SVTYPEs, which generally means that INSs from the input VCF only match when their coordinates are (nearly) the same as those in the BED file, even if the SVLENs between the potential matches differ. If the reciprocal overlap parameter is used, SVLENs for INSs are then used to better refine the matching INSs though differences in SVLENs may still exist between resulting INS matches. Other even more complex SVTYPEs, including copy number variants (CNVs), may require more specialized attention.

## Results

SVAFotate offers AF-related annotations that can greatly reduce the number of putative SVs for review in different genomic analyses. We demonstrate SVAFotate’s functionality by providing example analyses that categorize the rarity of SV calls based on their associated AFs, as well as illustrate how SVAFotate annotations can be used to effectively filter SVs. We also describe optional SVAFotate parameters that generate additional annotations to further enhance SV filtering and prioritization. Finally, we provide recommendations for applying SVAFotate in rare disease studies.

### Defining the rarity of SVs derived from CEPH families

We first highlight the utility of SVAFotate in classifying SVs based on their associated population frequency by creating SVAFotate annotations for SV calls derived from 603 individuals belonging to 34 multigenerational CEPH families (26). SVs were called for each individual and then merged into a single VCF using Smoove with the GRCh38 reference genome and parameterized as recommended in the tool’s documentation (https://github.com/brentp/smoove). The resulting VCF reported nearly 40,000 total SV calls across all individuals with a total of 21,106 deletions, 7,021 duplications, 686 inversions, and 10,702 unclassified breakend (BND) calls. SVAFotate annotations were then added using the provisional GRCh38 BED file corresponding to SV calls from the CCDG, gnomAD, and 1000G SV datasets while also requiring a reciprocal overlap of 80% (`-f 0.8)`. This annotation was accomplished with the following command:

~~~
$ svafotate annotate --vcf CEPH.smoove.vcf.gz --out svafotate.vcf -f 0.8 -b SVAFotate_core_SV_popAFs.GRCh38.bed.gz
~~~

Focusing on the deletion, duplication, and inversion calls and using the `Max_AF` annotation provided by SVAFotate, each SV was classified as *Common* (`Max_AF >= 0.05`), *LowFreq* (`0.05 > Max_AF >= 0.01`), *Rare* (`Max_AF < 0.01`), or *Unique* (`Max_AF = 0.0`). We find that just under half of all these SVs are unique or private to this CEPH dataset when compared to those reported by CCDG, gnomAD, and 1000G (**Figure 3a**).

**Figure 3.**
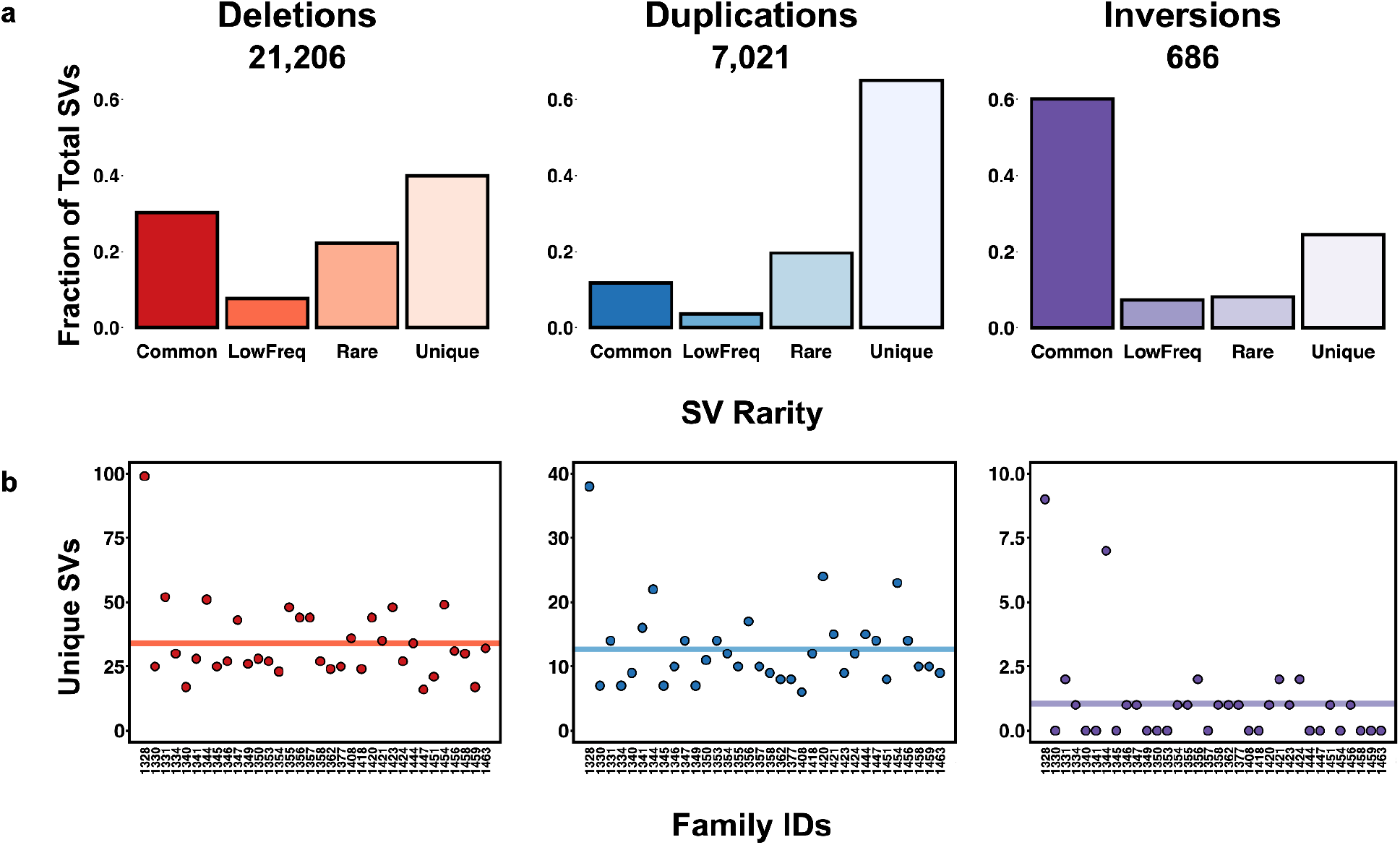
Frequency of CEPH SVs. **a**. Barplots representing the fraction of CEPH derived SVs per SVTYPE (deletions, duplications, and inversions) that are classified as *Common* (`Max_AF >= 0.05`), *LowFreq* (`0.05 > Max_AF >= 0.01`), *Rare* (`Max_AF < 0.01`), or *Unique* (`Max_AF = 0.0`). **b**. The total number of *Unique* SVs identified per SVTYPE (deletions, duplications, and inversions) that are CEPH family-specific with the mean indicated as a solid, colored line.

Using this classification we identified family-specific SVs that are labeled as *Unique* and only observed in a single CEPH family (**Figure 3b**). On average, each CEPH family harbors roughly 16 SV events that are private to that family and have no matches with any SVs of the same SVTYPE from CCDG, gnomAD, or 1000G. Family 1328 exhibits a higher number of unique SV events than any other family (146 total unique SVs), but is also an outlier with regards to the family size (83 total individuals compared to the median of 14.5 individuals per CEPH family). CEPH individuals are considered healthy with no clinically reported phenotypes so at this time these unique SVs are not considered as candidate variants for any specific condition. However, this analysis demonstrates how SVAFotate may assist in identifying rare SVs that are unique to specific families and may contribute to observed phenotypes, including rare diseases.

### Using SVAFotate annotations to filter SVs in neonatal ICU cases

A primary motivation for developing SVAFotate was to enable filtering of SV calls using known population AFs. We further demonstrate the use of SVAFotate annotations by applying them to rare disease cases from the Utah NeoSeq Project (27), a rapid WGS protocol to provide genetic diagnoses for critically ill infants in the University of Utah Hospital neonatal intensive care unit. Since the inception of the Utah NeoSeq Project, SVAFotate has been used in the SV analysis and prioritization pipeline. Here, we summarize the filtering of SVs from 22 NeoSeq cases, 19 of which are trios (proband and both parents) and 3 are duos (proband and a single parent).

SV calling was performed using the recommended parameters for both Smoove and Manta (28), using the GRCh38 reference genome, resulting in two SV-specific VCFs for each NeoSeq case. Smoove VCFs featured deletions, duplications, inversions, and BNDs while Manta VCFs included deletions, duplications, insertions, and BNDs. SVAFotate annotations were then added to each VCF with a required 80% reciprocal overlap. This was done using the provisional GRCh38 BED file corresponding to SV calls from the CCDG, gnomAD, and 1000G datasets. An additional SVAFotate annotated VCF was also generated using the same parameters while replacing the provisional BED file with a custom NeoSeq-specific GRCh38 BED file. This custom BED file was made using the same SV calls from the CCDG, gnomAD, and 1000G datasets, but also features SV calls and their AF related data from the aforementioned CEPH VCF as well as both Smoove and Manta calls derived from the 1000G samples. These additional SV calls were added to the custom BED file because they correspond to SVs derived from the same SV callers that are employed by the NeoSeq pipeline (Smoove and Manta). The inclusion of these additional calls should enable the identification of SVs that are more prone to being identified by these specific SV detection tools, and thus may be more representative of technical artifacts rather than true SV events.

For each VCF, the total number of SVs where the proband was called as heterozygous or homozygous for the alternate allele was determined, and we then counted how many of these SVs would be retained after imposing an AF filter that ranged between `0` and `1` using the `Max_AF` annotation provided by SVAFotate. For each AF cutoff value, the fraction of filtered SVs was determined by subtracting from one the number of retained SVs divided by the total number of variant SVs found in the proband. As expected, the fraction of SVs that are filtered increases as the AF cutoff is lowered (**Figure 4**).

**Figure 4.**
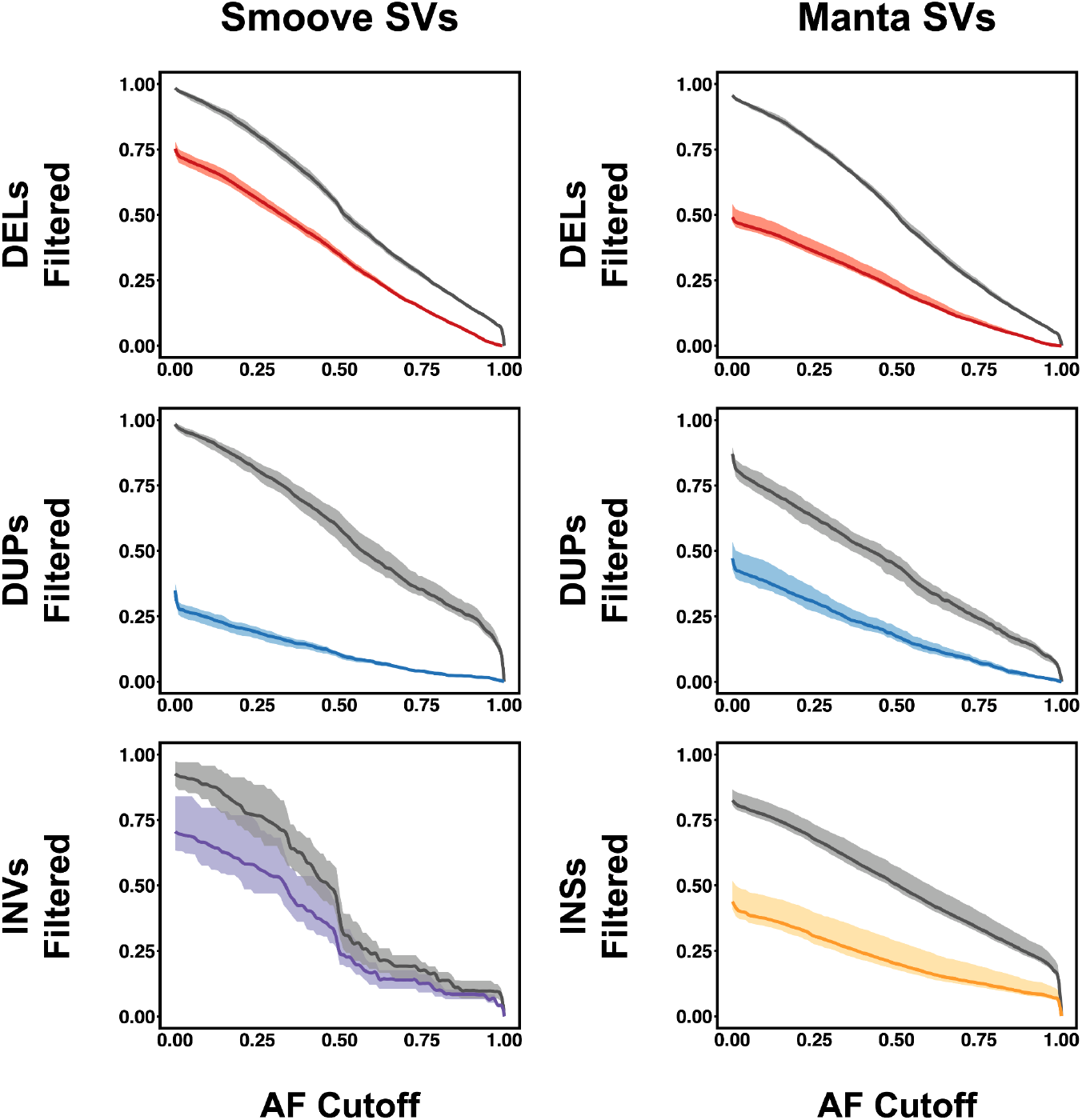
Filtering of NeoSeq SVs using AF cutoffs. The fraction of NeoSeq proband SV calls, per SVTYPE, that are filtered by using the `Max_AF` annotation added by SVAFotate and a range of AF cutoff values. SVTYPEs are abbreviated as follows: deletions (DELs), duplications (DUPs), inversions (INVs), and insertions (INSs). Plots on the left are SV calls derived from Smoove while the plots on the right are from Manta. Lines that are colored represent the resulting filtered SVs using the `Max_AF` annotation generated using the provisional BED file, while the gray lines represent the filtered SVs using the `Max_AF` annotation created by the custom NeoSeq BED file. Each line has the maximum and minimum amount of filtered SVs observed across all 22 NeoSeq cases analyzed plotted as a shadow surrounding the line.

For example, using an AF cutoff of `0.01` and the SVAFotate annotations added using the provisional BED file with SVs pertaining to only the CCDG, gnomAD, and 1000G datasets, we observe that nearly 60% of NeoSeq deletions, duplications, and inversions called by Smoove are filtered while more than 40% of Manta derived deletions, duplications, and insertions are removed. However, the same analysis using the custom NeoSeq BED file as an input when generating SVAFotate annotations results in over 95% and 85% of the same Smoove and Manta NeoSeq SV calls being filtered. These findings suggest an appreciable number of the SVs in the Smoove and Manta VCFs are likely the result of the SV detection softwares used rather than true SVs. Alternatively, these additionally filtered calls using the custom NeoSeq BED file may represent problematic SVs that were identified by the CCDG, gnomAD, and 1000G efforts, but failed to meet required standards and were subsequently removed in these study cohorts. Altogether, these results highlight the value of adding additional SV datasets to the provisional BED file to enable more comprehensive SV filtering.

### SVAFotate Best Practices

While SVAFotate’s default settings create a foundation of annotations that facilitate SV interpretation, we highlight additional options that provide more detailed annotations enabling deeper analyses and variant prioritization (**Table 1**).

**Table 1.** SVAFotate Parameters and Options. Required SVAFotate parameters are listed followed by optional SVAFotate parameters with all arguments and descriptions for each option included. Recommended optional parameters are listed in bold with accompanying recommendations, as applicable. For full descriptions and details, please refer to the SVAFotate repo: https://github.com/fakedrtom/SVAFotate

| Required SVAFotate Parameters | Argument | Description |
| --- | --- | --- |
| Input VCF | -v | The input VCF to be annotated by SVAFotate |
| Input BED | -b | The input BED of reported SVs to use for annotations |
| Output VCF | -o | The name for the resulting, SVAFotate annotated VCF |
| Optional SVAFotate Parameters | Argument | Description |
| Reciprocal overlap | -f | The minimum amount of reciprocal overlap required for the matching of SVs (recommended 0.8 or higher) |
| Extra annotations | -a | **Adds annotations corresponding to all extra annotations (best, full, mf, mis, pops)**<br>all: Adds all available information from the BED file for all matches<br>**best: adds the "best" match for each including source from the BED file**<br>mf: adds male and female specific data<br>**mis: Adds "mismatches" for matches with differing SVTYPES**<br>pop: Adds populations specific (AFR, AMR, EAS, EUR, OTH, and SAS) data, if available |
| **Observed SV coverage** | **-c** | **Creates annotation reflecting the amount of a given SV in the input VCF has been observed with the same SVTYPE from SVs in the BED file** |
| Unique SV regions | -u | Creates additional output file containing "unique" regions of a given SV that are not observed to have any overlaps with SVs in the BED file |
| SV size limit | -l | **Maximize size of SVs from the BED file to include alongside -c and -u parameters (recommended 1,000,000)** |
| Targets BED file | -t | **Identify overlaps with particular regions of interest reported in an additional BED file** |
| Sources to annotate | -s | Restrict matching of SVs to only those from specified sources in the BED file |
| Use CI boundaries | -ci | Adjust the genomic breakpoints in the input VCF by the reported inner or outer confidence intervals (CIPOS, CIEND) |
|  | -ci95 | Adjust the genomic breakpoints in the input VCF by the reported 95% inner or outer confidence intervals (CIPOS95, CIEND95) |
| Change SV size | -e | Increase the size of the genomic breakpoints in the input VCF |
|  | -r | Reduce the size of the genomic breakpoints in the input VCF |
| Output File Type | -O | Specify the output file type as VCF (vcf), compressed VCF (vcf.gz), BCF (bcf), or compressed BCF (bcf.gz) |
| CPU count | --cpu | The number of CPUs to use for multi-threading |

Based on use of SVAFotate in the previously mentioned Utah NeoSeq Project, which provides WGS of infants in the neonatal intensive care unit at the University of Utah, we describe in greater detail several of these optional settings and provide “best practices” for using SVAFotate towards the filtering and prioritization of SVs, particularly in rare disease analysis.

#### Reciprocal overlap fraction (-f)

Ideally, when identifying matches for the same structural variants in the input VCF and those found in the BED file of known SVs, identical genomic coordinates would be shared. However, given a multitude of variables involved in the calling of SVs, some discrepancies in genomic breakpoints are expected even for the same event present in multiple datasets. SVAFotate recognizes this uncertainty, and while it will identify any overlapping SV loci, properly matching the same SV events between datasets often requires greater similarity in the amount of overlap that exists between potential matches. By requiring a reciprocal overlap fraction, SVAFotate is better enabled to make more precise matches and thereby its allele frequency annotations are more representative of the measurements provided by the input BED file. This option requires that the overlap created by each SV in a potential match meets or exceeds a specified fraction of the total size of the SVs (**Figure 5a**). Based on observations from comparing the SVs from the CCDG, gnomAD, and 1000G datasets with one another (**Supplementary Figure 2**), a reciprocal overlap fraction of at least 80% is appropriate while still conservatively allowing some differences between potential matches.

**Figure 5.**
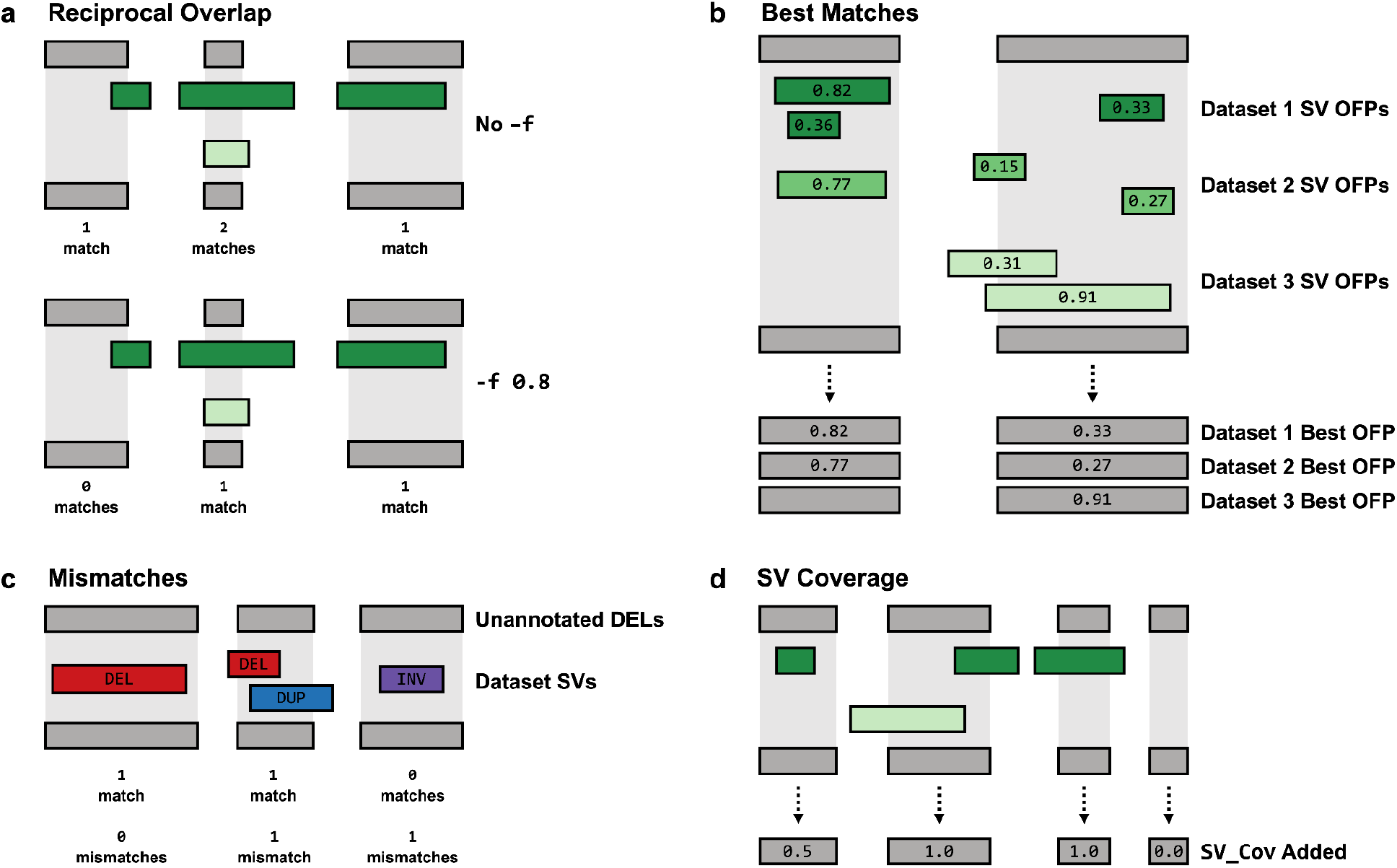
Recommended SVAFotate Parameters. Each plot illustrates SVs from the input VCF as gray rectangles with colored rectangles representing SVs from various datasets, such as CCDG, gnomAD, or 1000G. In all examples, the SVs depicted by gray and green rectangles are of the same SVTYPE. **a**. Requiring a reciprocal overlap with the -f parameter specifies that SVs being compared to one another must each have an overlap that meets a minimum fraction of the total size of the SV in order to be counted as a match and saved for future annotations by SVAFotate. On the top, the -f parameter is not being used and any overlap, regardless of size if being counted as a match, while on the bottom, -f is being used with a value of `0.8` which reduces the number of matches to those with greater overlap similarity. **b**. The OFPs for potential matches are calculated and listed as labels on each of the colored rectangles, representing SVs from three different datasets. The “best” match is determined by the match with the highest OFP value and metrics specific to that best match are saved and used for subsequent SVAFotate best annotations. If no match exists as illustrated for the SV on the left for Dataset 3, no best annotations are added for that dataset. **c**. Gray rectangles represent deletions from the input VCF and colored rectangles represent different SVTYPEs, specifically deletions (red), duplications (blue), and inversions (purple). Matches are defined as SVs of the same SVTYPE that overlap one another while mismatches are SVs of differing SVTYPEs that share an overlap. **d**. For each SV from the input VCF, all overlaps are saved and used to determine how much of the total SV region has also been observed in the datasets which is then reported as the `SV_Cov` annotation.

#### Extra annotations; best matches (-a best)

Any given SV may have multiple potential matches in the reference datasets, with each exhibiting differing genomic coordinates and AF metrics. While the core functionality of SVAFotate will return the maximum values from all of these potential matches, it is possible to also create annotations that reflect the “best” match, per source, based on genomic coordinate similarity. This is accomplished by computing an Overlap Fraction Product (OFP) between the input SV and all matching SVs from each source in the BED file. The OFP reflects the genomic similarity between matches by measuring the amount of overlap that is shared between the matches, calculating the fraction of each SV that the overlap covers, and then multiplying these fractions together (**Supplementary Figure 3**). Resulting OFP scores will have a range between `0.0` and `1.0` where high OFP scores reflect matching SVs that are more identical in terms of both their genomic sizes and the amount of overlap they share. Low OFP scores suggest a larger disparity in genomic sizes between matching SVs or a low amount of shared overlap between them (or both a discrepancy in sizes and low overlap). By reviewing the OFP scores from all matching SVs, a best match for each included source is determined by the matching SV with the highest OFP (**Figure 5b**). In the event of multiple matches sharing the same OFP score, the matching SV with the highest reported AF is returned as the best matching SV. Regardless of whether a reciprocal overlap is requested at the command line or not, determining the best match will consider all genomic overlaps. Thus, if any overlap exists for a given source reported in the BED file, regardless of its size, this will be reflected in the resulting best annotations. We determined the best match for all SVs by comparing the CCDG, gnomAD, and 1000G datasets to one another and observed largely bimodal distributions for all comparisons, suggesting that most best matches are either rather poor or quite precise (**Supplementary Figure 4**).

Once the best match is determined, multiple best annotations, including the OFP score for the best match, are saved to be added to the resulting output VCF. These can be used to help corroborate the values observed in the default annotations, but also provide more specificity with regards to which SVs from the input BED file are resulting in the SV matches. Furthermore, the best match allows for checking the precision of the match via the OFP annotation and the rarity or uniqueness of the input SV can be ascertained if no best matches exist.

#### Extra annotations; mismatches (-a mis)

By definition, SVAFotate requires matching SVs to share the same variant type (SVTYPE; e.g., deletion, duplication, etc.), but is also equipped to create annotations based on overlapping SVs with differing SVTYPEs when this option is used. SVAFotate refers to such overlaps as “mismatches” and are otherwise treated the same as traditionally matching SVs (**Figure 5c**). Mismatch annotations that are added include the differing SVTYPEs identified in the mismatches and also a series of best mismatch annotations similar to those created by the previously discussed best parameter in both methodology and content. Similar to the best annotations, mismatches are also determined for each source included in the input BED file.

Mismatches can reveal additional information for a given genomic region. For example, copy number variable loci often give rise to some individuals harboring deletions with others having duplications. On the other hand, such regions may indicate technically problematic regions for many SV detection tools. Mismatches may also represent situations where the same SV has been categorized differently between SV callers or datasets. For example, while some callers may identify and label an event as an insertion, others may classify the same event as a duplication. The mismatches annotations can be helpful in identifying such instances. Lastly, interpreting the potential phenotypic consequence of a rare or unique SV event might be influenced by the presence of a common SV of another SVTYPE that the mismatch annotations can reveal.

#### Observed SV Coverage (-c)

While overlapping SVs may represent distinct alleles, it may be informative to the interpretation of the possible pathogenicity of an SV if it occurs in the same genomic region as other SVs. The observed SV coverage (`SV_Cov`) annotation can help reveal these occurrences by describing the proportion of a given SV that overlaps SVs of the same SVTYPE from the input BED file. All overlaps with the same SVTYPE from all included sources, regardless of size, are considered when calculating this fraction and is reported as a number ranging from `0.0` to `1.0`, where high scores reflect a larger proportion of the SV being “covered” or observed to overlap with known SVs (**Figure 5d**). Additionally, similar coverage annotations that are source-specific are also added. This parameter also expects an AF cutoff value that will omit any SVs from the input BED file that fall below this AF threshold from the calculation of `SV_Cov`.

If any SVs in the BED file are exceedingly large, they may overwhelm the observed SV coverage, reducing its utility. For example, CCDG reports an exceptionally rare deletion that is over 61Mb in size. This event is likely to overlap with many putative deletions that likely represent distinct SV events. Considering this may obscure meaningful coverage annotations and interpretations, a recommended additional parameter to use alongside the observed SV coverage option is provided with the SV size limit parameter (**-l**). This limits the size of SVs from the input BED file to include when computing the observed SV coverage annotation. Given that many excessively large SVs defined by many variant callers can be spurious, we recommend setting the size limit to a megabase (1,000,000 bp) which will not include any SVs over that size when computing the observed SV coverage.

#### Targets BED File (-t)

Especially in the case of rare disease analysis, there may be particular genomic regions (e.g., genes known to be associated with the phenotype) where an overlap with any reported SV event would be of interest. Using this option and supplying a simple BED file consisting of chromosome, start and end coordinates, and a column featuring a region identifier (such as a gene name) will create a `Target_Overlaps` annotation that lists the supplied region identifier for all overlaps between the SV and the regions in this BED file.

Regions of interest may include any genomic features such as candidate genes, specific exons, promoters, enhancers, or any other set of user-determined genomic coordinates.

The following command details the use of these recommend options when using SVAFotate with the previously described and provided BED file and results in a SVAFotate annotated VCF named `svafotate.vcf`:

~~~
$ svafotate annotate --vcf input.vcf.gz --out svafotate.vcf -b
SVAFotate_core_SV_popAFs.GRCh38.bed.gz -f 0.8 -a best mis -c 0 -l 1000000 -t targets.bed
~~~

These SVAFotate annotations aid rare disease analysis by facilitating the identification of variants that are rare or unique to the affected individuals. As previously demonstrated, the default `Max_AF` annotation enables the classification of SVs based on their apparent rarity and therefore serves as the primary filter for categorizing SVs as rare or unique (`Max_AF < 0.01` and `Max_AF = 0.0`, respectively). However, there are important caveats to consider when reviewing any SVs that have passed this initial AF filter.

First, while a single SV event may be regarded as rare, it may reside in a locus that has been observed to harbor more commonly occurring SVs of the same SVTYPE and is thus less likely to be causative or otherwise qualify as a candidate variant. The `SV_Cov` annotation can help identify such occurrences and may serve as an additional filter. Any rare SV with an `SV_Cov` less than `1.0` contains some amount of genomic space that has not been previously observed as variable for that SVTYPE and an `SV_Cov` of `0.0` may be used to help identify rare SVs that are more independent of other variants. Second, reviewing possible mismatch annotations can also help determine how much other structural variation occurs within or near the same genomic region. This may affect the possible functional interpretation of a rare SV, especially if those mismatches are more common in the included datasets. Mismatch annotations also include OFP scores which may help prioritize or exclude mismatches under review. Lastly, best match annotations, or lack thereof, can also aid in identifying rare SVs that are in fact unique, with no known matches of any kind to the included dataset SVs. If no best matches exist for a given SV, then no overlaps of any kind with other SVs were found, meaning the SV is more likely to be unique. However, if any overlaps exist, these will be reported by the best annotations, where the exactness of the overlap is detailed in the best OFP annotation. These variants can possibly be disregarded if the OFP is not suggestive of a good match (OFP < `0.8`). Even SVs with best annotations may still represent unique SVs provided the best OFP scores are low. Additionally, best match annotations can help identify the source and exactness of the matches that contribute to the AF reported by the `Max_AF` annotation. In this manner, the use of the Max_AF, SV_Cov, and the mismatch and best annotations provided by SVAFotate work in combination to better determine the rarity and uniqueness of SVs within a VCF. In most cases, filtering using these annotations can be performed sequentially or simultaneously using available command line tools or custom scripts.

Together, these suggestions are sensible starting points for the filtering of SVs, but depending on the datasets used and the context of the SV analysis, they are all adjustable. SVs that do not meet these requirements may still be considered for further review. Once rare and unique SVs have been determined using SVAFotate, further filtering can be done based on expected inheritance patterns (*de novo* or dominant versus recessive) and other genetic features that are commonly included via additional annotation tools. If *a priori* there are genomic regions of interest, such as candidate genes, the `Target_Overlaps` annotation that can be included by SVAFotate can facilitate the identification of SVs that overlap these features and may be used as another filter. Otherwise, other gene annotation tools can enable the identification of SVs that overlap gene coding or other genic regions. Altogether, SVAFotate annotations are complementary to other genomic annotations and are meant to be used together in SV filtering and analysis.

## Conclusions

SVAFotate is ideally suited to combining and converting data from multiple SV datasets that contain population AF information into discrete annotations that can then be used for categorizing the rarity of SV calls and filtering them based on AF-related expectations. This is primarily accomplished by identifying matches based on user-defined genomic overlaps between queried SV calls in the VCF format and known SVs with associated AF features that are provided as a BED file. Population AF information from these matches is saved and summarized by SVAFotate as new and easily filterable annotations in a resulting output VCF file. SVAFotate is relatively fast, depending on the size of the input VCF and BED files, with most runtimes completing in less than 10 minutes for moderately sized VCF files (25,000 variant entries or less) and the provided BED files.

Rare disease analysis benefits from the ability to determine the frequency of variants in general or specific populations. Multiple methods and datasets are available to do this for common variant types, like SNVs and INDELs, but SVs can be more problematic in these types of analyses. SVAFotate enables the annotation of SVs with population frequencies and other similar data obtained from previously identified SVs. As SV datasets continue to grow and become available, especially with accompanying population-level measurements, these can be added to the expected SVAFotate input BED file to provide more comprehensive SV annotations. Furthermore, SVAFotate is designed to allow for the addition of specific or custom types of SV datasets, such as those generated using specific variant callers, to allow for precise SV matching and subsequent filtering. This positions SVAFotate as a valuable resource for inclusion in current and future SV analyses.

While SVAFotate is not clinically diagnostic and does not rank SV calls itself, the annotations it creates provides the information necessary to rapidly sort through SV calls and determine their apparent rarity with regards to known SV datasets. Filtering based on these AF metrics can substantially reduce a given SV call set and thus, effectively prioritize SV calls, especially in the context of rare disease analysis. Combining SVAFotate annotations with other common genetic features, such as various gene annotations or other SV prioritization tools, can further refine this list of SV calls resulting in manageable lists of variants for manual review and verification.

SVAFotate is an open-source software package and it is freely available. Source code and further documentation can be found at: https://github.com/fakedrtom/SVAFotate.

## Availability and requirements

Project name: SVAFotate

Project home page: https://github.com/fakedrtom/SVAFotate

Operating system(s): Platform independent

Programming language: Python

Other requirements: None

License: MIT

Any restrictions to use by non-academics: None

## List of abbreviations

AF: Allele Frequency
SV: Structural Variant
SNV: Single Nucleotide Variant
INDEL: Insertion-Deletion
WGS: Whole-Genome Sequencing
DEL: Deletion
DUP: Duplication
INV: Inversion
INS: Insertion
BND: Unclassified Breakend
OFP: Overlap Fraction Product

## Declarations

### Ethics approval and consent to participate

Informed consent was obtained from the CEPH individuals, and the University of Utah Institutional Review Board approved the study (University of Utah IRB #80145). Written informed consent was obtained from all participants, including written consent from the parents or legal guardians of any participant under the age of 16, in the Utah NeoSeq project which was approved by the University of Utah Institutional Review Board (University of Utah IRB #00125940).

### Consent for publication

Not applicable.

### Availability of data and materials

Provisional input BED files corresponding to SV calls from the CCDG, gnomAD, and 1000G datasets are provided for the GRCh37 and GRCh38 human genome reference builds. Additionally, data from the probands of Utah NeoSeq cases that correspond to SV calls generated by Smoove and Manta, their corresponding genotypes, SVTYPEs, and Max_AF annotations from SVAFotate using both the provisional BED and custom NeoSeq BED files are also provided and can be found here: https://doi.org/10.5281/zenodo.6506679

The analyses included utilize CEPH data that has been previously published. Aligned sequencing reads (in CRAM format) pertaining to CEPH samples are available at the SRA and dbGaP under controlled access, with accession phs001872.v1.p1.

### Authors’ contributions

TJN designed and developed SVAFotate. TJN and MJC provided code improvement. TJN wrote the manuscript. All authors read and approved the final manuscript.

## Acknowledgements

The authors wish to thank the CCDG, gnomAD, and 1000G groups for making their SV datasets publicly available. The authors also thank Laurel Hiatt, Stephanie Kravitz, and Matt Velinder for valuable feedback and suggestions regarding the manuscript.

