## supplementary_materials for "Annotation of structural variants with reported allele frequencies and related metrics from multiple datasets using SVAFotate"

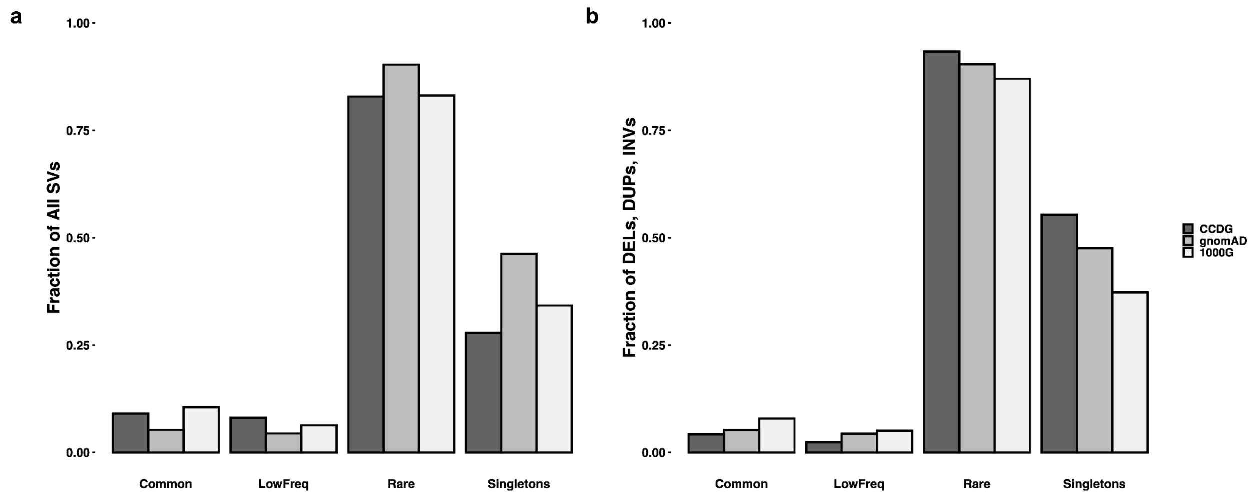

**Supplementary Figure 1. Reported SV Rarities by CCDG, gnomAD, and 1000G**

The fraction of SVs reported by CCDG, gnomAD, and 1000G found to be “Common” (AF >= 0.05), “LowFreq” (0.05 > AF >= 0.01), and “Rare” (AF < 0.01). Additionally, “Singletons” where allele count in genotypes is 1 (AC = 1), which are inherently a subset of the “Rare” SVs, are also plotted. **a.** Features all SVs reported by each dataset where on average, approximately 8% are “Common”, 6% are “LowFreq”, and 85% are “Rare” with 36% of all SVs being “Singletons” . **b.** Limits the SVs to SVTYPEs that the datasets have in common (deletions, duplications, and inversions) where on average approximately 6% are “Common”, 4% are “LowFreq”, and 90% are “Rare” with 47% of all SVs being “Singletons”.

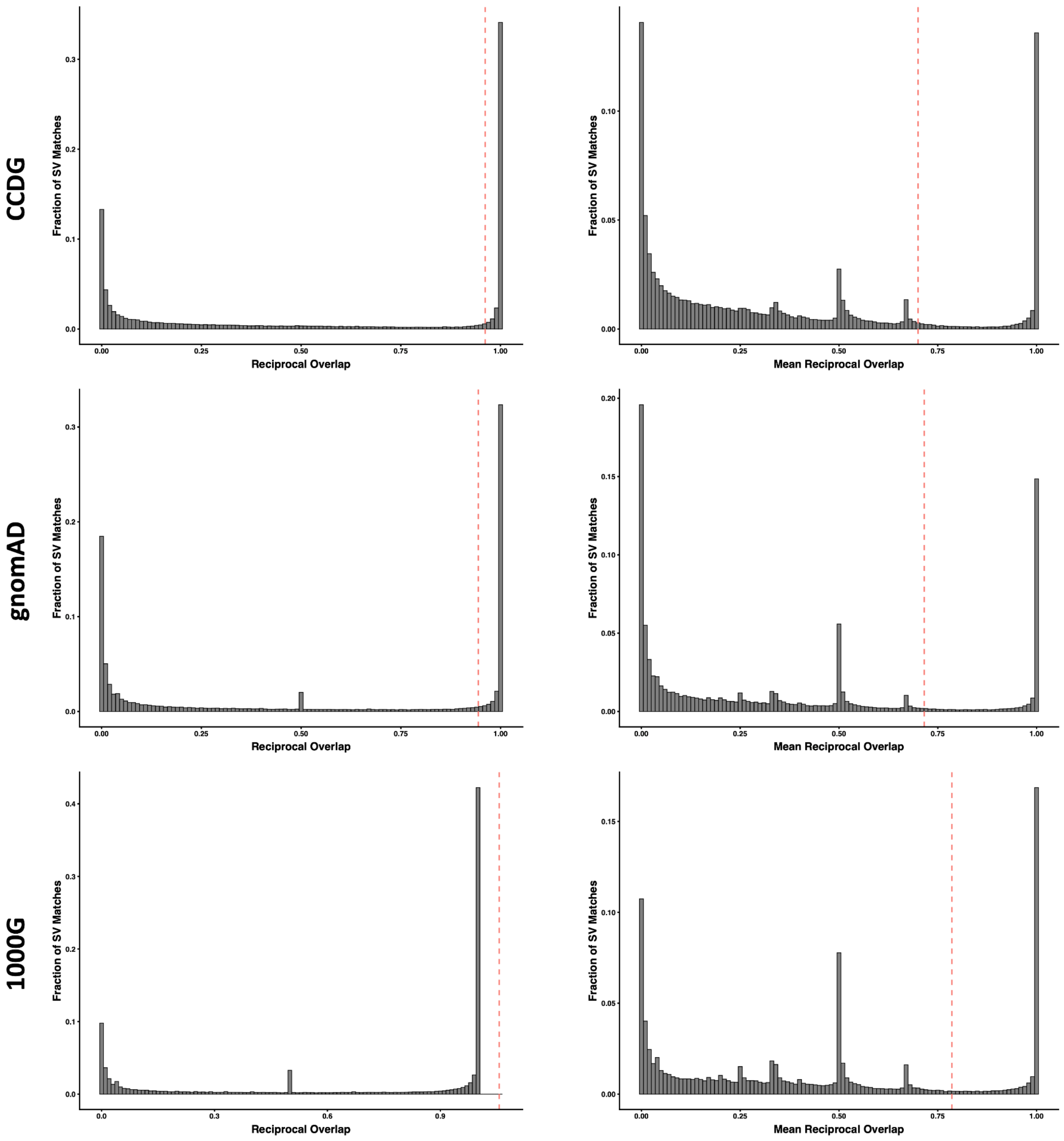

**Supplementary Figure 2. Distributions of the Reciprocal Overlap Identified Across Matches from Dataset Comparisons**

For the CCDG, gnomAD, and 1000G datasets, distributions of reciprocal overlaps observed between all possible matches between the datasets is plotted for each dataset with one standard deviation beyond the mean indicated as a red dashed line. On the left, distributions reflecting only the highest observed reciprocal overlap for each given SV in a dataset is plotted while on the right, distributions representing the mean observed reciprocal overlap for all matches to a given SV in a dataset is shown.

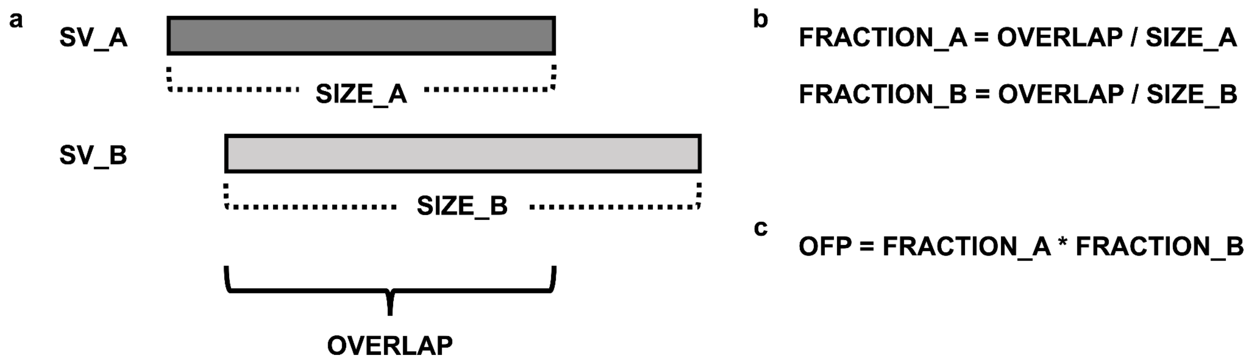

**Supplementary Figure 3. Calculating the Overlap Fraction Product (OFP)**

**a.** For each overlapping pair of SVs, the amount of overlap is determined. **b.** Using the overlap, the fraction of each SV that is found to overlap with the other is calculated. **c.** These fractions are then multiplied to create the Overlap Fraction Product (OFP).

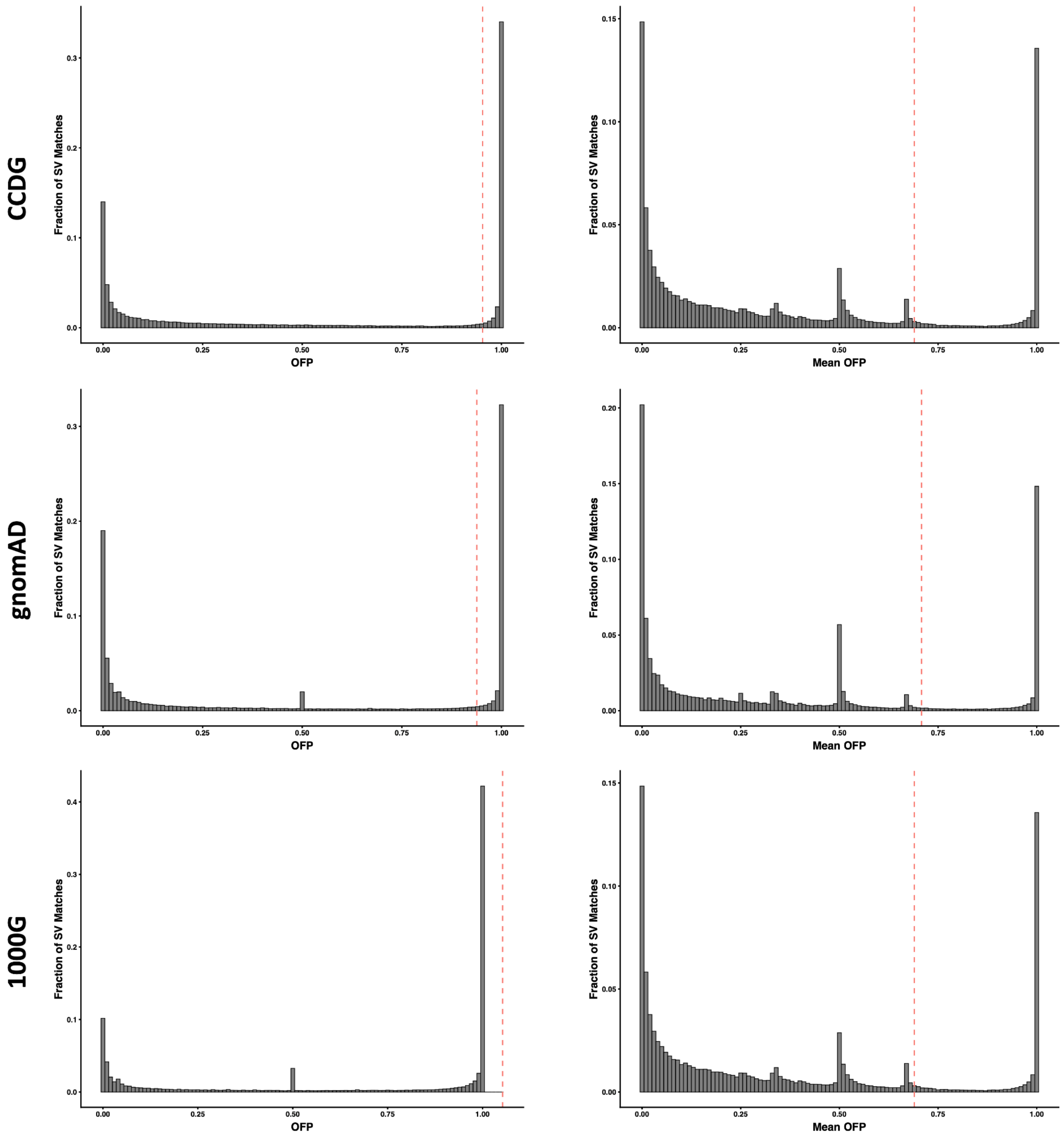

### Supplementary Figure 4. Distributions of OFPs Across Dataset Comparisons

For the CCDG, gnomAD, and 1000G datasets, distributions of OFPs observed between all possible matches between the datasets is plotted for each dataset with one standard deviation beyond the mean indicated as a red dashed line. On the left, distributions reflecting only the highest or best observed OFP for each given SV in a dataset is plotted while on the right distributions representing the mean OFP for all matches to a given SV in a dataset is shown.

### Supplementary Table 1. Expected Columns for Input BED File

| Expected Column | Description |
| --- | --- |

| | |
|---|---|
| CHROM | Chromosome |
| START | Start position of SV |
| END | End position of SV |
| SVLEN | Length of SV |
| SVTYPE | Type of SV |
| SOURCE | Data source that reports the SV |
| SV_ID | Unique identifier for the SV |
| AF | Allele frequency |
| HomRef | Count of individuals with homozygous for the reference allele genotype |
| Het | Count of individuals with heterozygous genotype |
| HomAlt | Count of individuals with homozygous for the alternate allele genotype |
| Male_AF | Allele frequency of male samples |
| Male_HomRef | Count of male individuals with homozygous for the reference allele genotype |
| Male_Het | Count of male individuals with heterozygous genotype |
| Male_HomAlt | Count of male individuals with homozygous for the alternate allele genotype |
| Male_HemiAlt | Count of male individuals with hemizygous genotype |
| Male_HemiAF | Hemizgous allele frequency |
| Female_AF | Allele frequency of female samples |
| Female_HomRef | Count of female individuals with homozygous for the reference allele genotype |
| Female_Het | Count of female individuals with heterozygous genotype |
| Female_HomAlt | Count of female individuals with homozygous for the alternate allele genotype |
| AFR_AF | Allele frequency of AFR samples |
| AFR_HomRef | Count of AFR individuals with homozygous for the reference allele genotype |
| AFR_Het | Count of AFR individuals with heterozygous genotype |
| AFR_HomAlt | Count of AFR individuals with homozygous for the alternate allele genotype |
| AFR_Male_AF | Allele frequency of AFR male samples |
| AFR_Male_HomRef | Count of AFR male individuals with homozygous for the reference allele genotype |
| AFR_Male_Het | Count of AFR male individuals with heterozygous genotype |
| AFR_Male_HomAlt | Count of AFR male individuals with homozygous for the alternate allele genotype |
| AFR_Male_HemiAlt | Count of AFR male individuals with hemizygous genotype |
| AFR_Male_HemiAF | Hemizgous allele frequency of AFR samples |
| AFR_Female_AF | Allele frequency of AFR female samples |
| AFR_Female_HomRef | Count of AFR female individuals with homozygous for the reference allele genotype |

| | |
|---|---|
| AFR_Female_Het | Count of AFR female individuals with heterozygous genotype |
| AFR_Female_HomAlt | Count of AFR female individuals with homozygous for the alternate allele genotype |
| AMR_AF | Allele frequency of AMR samples |
| AMR_HomRef | Count of AMR individuals with homozygous for the reference allele genotype |
| AMR_Het | Count of AMR individuals with heterozygous genotype |
| AMR_HomAlt | Count of AMR individuals with homozygous for the alternate allele genotype |
| AMR_Male_AF | Allele frequency of AMR male samples |
| AMR_Male_HomRef | Count of AMR male individuals with homozygous for the reference allele genotype |
| AMR_Male_Het | Count of AMR male individuals with heterozygous genotype |
| AMR_Male_HomAlt | Count of AMR male individuals with homozygous for the alternate allele genotype |
| AMR_Male_HemiAlt | Count of AMR male individuals with hemizygous genotype |
| AMR_Male_HemiAF | Hemizgous allele frequency of AMR samples |
| AMR_Female_AF | Allele frequency of AMR female samples |
| AMR_Female_HomRef | Count of AMR female individuals with homozygous for the reference allele genotype |
| AMR_Female_Het | Count of AMR female individuals with heterozygous genotype |
| AMR_Female_HomAlt | Count of AMR female individuals with homozygous for the alternate allele genotype |
| EAS_AF | Allele frequency of EAS samples |
| EAS_HomRef | Count of EAS individuals with homozygous for the reference allele genotype |
| EAS_Het | Count of EAS individuals with heterozygous genotype |
| EAS_HomAlt | Count of EAS individuals with homozygous for the alternate allele genotype |
| EAS_Male_AF | Allele frequency of EAS male samples |
| EAS_Male_HomRef | Count of EAS male individuals with homozygous for the reference allele genotype |
| EAS_Male_Het | Count of EAS male individuals with heterozygous genotype |
| EAS_Male_HomAlt | Count of EAS male individuals with homozygous for the alternate allele genotype |
| EAS_Male_HemiAlt | Count of EAS male individuals with hemizygous genotype |
| EAS_Male_HemiAF | Hemizgous allele frequency of EAS samples |
| EAS_Female_AF | Allele frequency of EAS female samples |
| EAS_Female_HomRef | Count of EAS female individuals with homozygous for the reference allele genotype |
| EAS_Female_Het | Count of EAS female individuals with heterozygous genotype |
| EAS_Female_HomAlt | Count of EAS female individuals with homozygous for the alternate allele |

| | |
|---|---|
| | genotype |
| EUR_AF | Allele frequency of EUR samples |
| EUR_HomRef | Count of EUR individuals with homozygous for the reference allele genotype |
| EUR_Het | Count of EUR individuals with heterozygous genotype |
| EUR_HomAlt | Count of EUR individuals with homozygous for the alternate allele genotype |
| EUR_Male_AF | Allele frequency of EUR male samples |
| EUR_Male_HomRef | Count of EUR male individuals with homozygous for the reference allele genotype |
| EUR_Male_Het | Count of EUR male individuals with heterozygous genotype |
| EUR_Male_HomAlt | Count of EUR male individuals with homozygous for the alternate allele genotype |
| EUR_Male_HemiAlt | Count of EUR male individuals with hemizygous genotype |
| EUR_Male_HemiAF | Hemizgous allele frequency of EUR samples |
| EUR_Female_AF | Allele frequency of EUR female samples |
| EUR_Female_HomRef | Count of EUR female individuals with homozygous for the reference allele genotype |
| EUR_Female_Het | Count of EUR female individuals with heterozygous genotype |
| EUR_Female_HomAlt | Count of EUR female individuals with homozygous for the alternate allele genotype |
| OTH_AF | Allele frequency of OTH samples |
| OTH_HomRef | Count of OTH individuals with homozygous for the reference allele genotype |
| OTH_Het | Count of OTH individuals with heterozygous genotype |
| OTH_HomAlt | Count of OTH individuals with homozygous for the alternate allele genotype |
| OTH_Male_AF | Allele frequency of OTH male samples |
| OTH_Male_HomRef | Count of OTH male individuals with homozygous for the reference allele genotype |
| OTH_Male_Het | Count of OTH male individuals with heterozygous genotype |
| OTH_Male_HomAlt | Count of OTH male individuals with homozygous for the alternate allele genotype |
| OTH_Male_HemiAlt | Count of OTH male individuals with hemizygous genotype |
| OTH_Male_HemiAF | Hemizgous allele frequency of OTH samples |
| OTH_Female_AF | Allele frequency of OTH female samples |
| OTH_Female_HomRef | Count of OTH female individuals with homozygous for the reference allele genotype |
| OTH_Female_Het | Count of OTH female individuals with heterozygous genotype |
| OTH_Female_HomAlt | Count of OTH female individuals with homozygous for the alternate allele genotype |
| SAS_AF | Allele frequency of SAS samples |

| | |
|---|---|
| SAS_HomRef | Count of SAS individuals with homozygous for the reference allele genotype |
| SAS_Het | Count of SAS individuals with heterozygous genotype |
| SAS_HomAlt | Count of SAS individuals with homozygous for the alternate allele genotype |
| SAS_Male_AF | Allele frequency of SAS male samples |
| SAS_Male_HomRef | Count of SAS male individuals with homozygous for the reference allele genotype |
| SAS_Male_Het | Count of SAS male individuals with heterozygous genotype |
| SAS_Male_HomAlt | Count of SAS male individuals with homozygous for the alternate allele genotype |
| SAS_Male_HemiAlt | Count of SAS male individuals with hemizygous genotype |
| SAS_Male_HemiAF | Hemizgous allele frequency of SAS samples |
| SAS_Female_AF | Allele frequency of SAS female samples |
| SAS_Female_HomRef | Count of SAS female individuals with homozygous for the reference allele genotype |
| SAS_Female_Het | Count of SAS female individuals with heterozygous genotype |
| SAS_Female_HomAlt | Count of SAS female individuals with homozygous for the alternate allele genotype |
| PopMax_AF | The maximum AF across all populations |
| InPop | The number of populations that report the SV |

**Supplementary Table 2. SVAFotate Annotations**

| SVAFotate Parameter | Added Annotation | Description |
|---|---|---|
| default | Max_AF | The maximum AF from all matching SVs |
| | Max_Het | The maximum count of heterozygote genotypes from all matching SVs |
| | Max_HomAlt | The maximum count of homozygote alternate genotypes from all matching SVs |
| | Max_PopMax_AF | The maximum PopMax_AF from all matching SVs |
| -a best | Best_[data_source]_ID | The SV ID of the best matching SV for that data source |
| | Best_[data_source]_OFP | The OFP of the best matching SV for that data source |
| | Best_[data_source]_AF | The AF of the best matching SV for that data source |
| | Best_[data_source]_Het | The count of heterozygous genotypes for best matching SV for that data source |
| | Best_[data_source]_HomAlt | The count of homozygous alternate genotypes for best matching SV for that data source |
| | Best_[data_source]_PopMax_AF | The PopMax_AF of the best matching SV for that data source |
| -a mf | Max_Male_AF | The maximum male AF from all matching SVs |
| | Max_Male_Het | The maximum male count of heterozygous genotypes from all matching SVs |
| | Max_Male_HomAlt | The maximum male count of homozygous alternate genotypes from all matching SVs |
| | Max_Female_AF | The maximum female AF from all matching SVs |
| | Max_Female_Het | The maximum female count of heterozygous genotypes from all matching SVs |
| | Max_Female_HomAlt | The maximum female count of homozygous alternate genotypes from all matching SVs |
| -a mis | [data_source]_Mismatches | List of SV IDs from all overlapping SVs with a different SVTYPE for that data source |
| | [data_source]_Mismatches_Count | The number of mismatches identified for that data source |
| | [data_source]_Mismatches_SVTYPEs | The other SVTYPEs identified from overlapping SVs with a different SVTYPE for that data source |
| | Best_[data_source]_Mismatch_ID | The SV ID of the best mismatching SV for that data source |
| | Best_[data_source]_Mismatch_OFP | The OFP of the best mismatching SV for that data source |
| | Best_[data_source]_Mismatch_SVTYPE | The SVTYPE of the best mismatching SV for that data source |
| | Best_[data_source]_Mismatch_AF | The AF of the best mismatching SV for that data source |
| | Best_[data_source]_Mismatch_Het | The count of heterozygous genotypes for best mismatching SV for that data source |
| | Best_[data_source]_Mismatch_HomAlt | The count of homozygous alternate genotypes for best mismatching SV for that data source |
| -a pops | Max_[population]_AF | The maximum population specific AF from all matching SVs |
| | Max_[population]_Het | The maximum population specific count of heterozygote genotypes from all matching SVs |
| | Max_[population]_HomAlt | The maximum population specific count of homozygote alternate genotypes from all matching SVs |
| -c | SV_Cov | The fraction of the SV that has been observed with the same SVTYPE across all data sources |
| | [data_source]_SV_Cov | The fraction of the SV that has been observed with the same SVTYPE for that data source |
| -u | SV_Uniq | The number of unique regions found within the SV |
| -t | Target_Overlaps | The region identifier(s) for all overlaps between the SV and the regions of interest in the targets BED file |
